# A Machine Learning-optimized system for on demand, pulsatile, photo- and chemo-therapeutic treatment using near-infrared responsive MoS_2_-based microparticles in a breast cancer model

**DOI:** 10.1101/2023.04.16.536750

**Authors:** Maria Kanelli, Neelkanth M. Bardhan, Morteza Sarmadi, Shahad Alsaiari, William T. Rothwell, Apurva Pardeshi, Dominique C. De Fiesta, Howard Mak, Virginia Spanoudaki, Nicole Henning, Jooli Han, Angela M. Belcher, Robert S. Langer, Ana Jaklenec

## Abstract

Cancer therapy research is of high interest because of the persistence and mortality of the disease and the side effects of traditional therapeutic methods, while often multimodal treatments are necessary based on the patient’s needs. The development of less invasive modalities for recurring treatment cycles is thus of critical significance. Herein, a light-activatable microparticle system was developed for localized, pulsatile delivery of anticancer drugs with simultaneous thermal ablation, by applying controlled ON-OFF thermal cycles using near-infrared laser irradiation. The system is composed of poly(caprolactone) microparticles of 200 μm size with incorporated molybdenum disulfide (MoS_2_) nanosheets as the photothermal agent and hydrophilic doxorubicin or hydrophobic violacein, as model drugs. Upon irradiation the nanosheets heat up to ≥50 °C leading to polymer matrix melting and release of the drug. MoS_2_ nanosheets exhibit high photothermal conversion efficiency and allow for application of low power laser irradiation for the system activation. A Machine Learning algorithm was applied to acquire optimal laser operation conditions; 0.4 W/cm^2^ laser power at 808 nm, 3-cycle irradiation, for 3 cumulative minutes. In a mouse subcutaneous model of 4T1 triple-negative breast cancer, 25 microparticles were intratumorally administered and after 3-cycle laser treatment the system conferred synergistic phototherapeutic and chemotherapeutic effect. Our on-demand, pulsatile synergistic treatment resulted in increased median survival up to 40 days post start of treatment compared to untreated mice, with complete eradication of the tumors at the primary site. Such a system could have potential for patients in need of recurring cycles of treatment on subcutaneous tumors.

**GRAPHICAL ABSTRACT:** 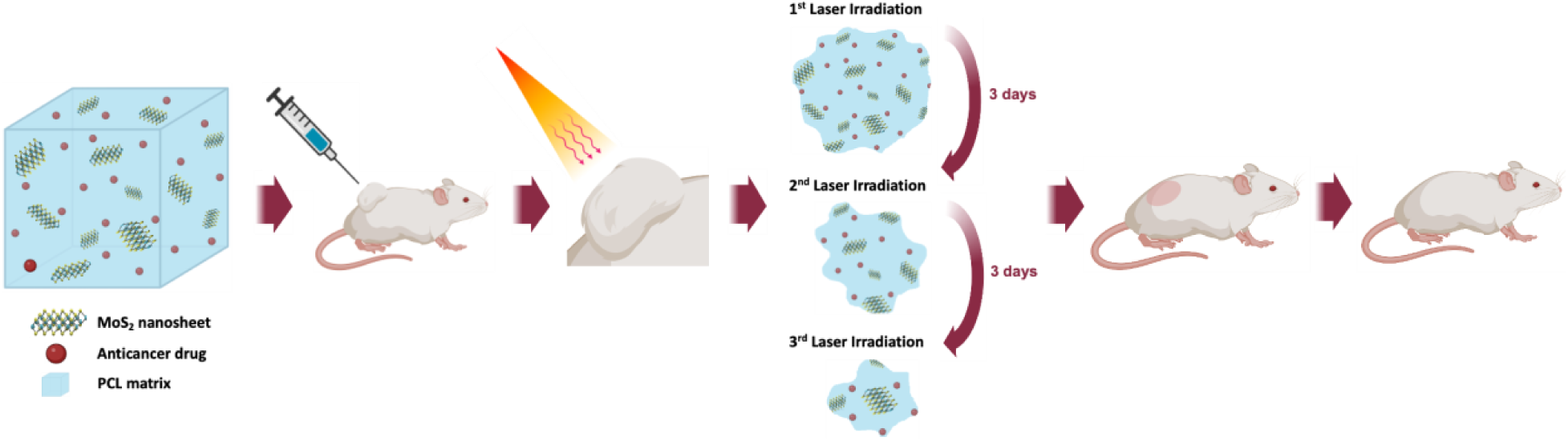

## INTRODUCTION

Cancer is one of the most prevalent causes of human demise throughout the world, while the main treatment routes involve surgery, radiotherapy and chemotherapy [1]. While great strides have been made towards cancer treatment over the past years, it continues to be a major health concern. Furthermore, many drugs are cytotoxic both to tumor and healthy cells and, thus, they present side effects such as hair loss, nausea and gastrointestinal tract lesions [2]. Therefore, extensive efforts have been devoted to searching for improved therapeutic approaches.

Since cancer is a complex disease, with high patient-to-patient variability in the genetic expression profile as well as the tumor microenvironment, the successful clinical management often requires a multi-modal treatment approach, using surgery, chemotherapy, radiation therapy, or immunotherapy in certain cases, to name a few. Specifically, in the case of metastatic breast cancer, many patients require multiple rounds of treatment, which can be physically and mentally challenging for the patient to endure. Thus, there is an unmet clinical need to develop less invasive, nontoxic, and locally targeted multi-modal therapies which can help alleviate the burden of successive rounds of chemotherapies and other cyclic treatment options and mitigate the side effects of systemic toxicity. Herein we propose a local therapeutic modality, with a controlled, pulsatile drug release mechanism. There have been several studies that harness the photothermal, photochemical or photomechanical interactions of light with tissues and cells. Two of the most commonly used modalities are photodynamic therapy (PDT) and photothermal therapy (PTT) [3]. In the former, light-activatable molecules, or “photosensitizers” are delivered to the tumor tissue, and upon irradiation with light, leads to activation resulting in the generation of free radicals or reactive oxygen species that are locally cytotoxic to the tumor microenvironment. In the latter, light irradiation is used to achieve heat generation, resulting in hyperthermia over threshold coagulation temperatures of ∼ 50-55 °C. Photothermal heating can also enhance the permeability of the cell membrane and vascular permeability, resulting in higher trafficking and intracellular uptake of anticancer drugs [4]. Typically, this is achieved through the administration of nanoparticle-based thermal enhancement agents which absorb preferentially in the near-infrared range of wavelengths, allowing for deeper penetration into the tissue for killing bulk tumors [5]. In recent years, the use of near-infrared (NIR: 700 - 1700 nm) optical techniques for functionalized nanomaterials has emerged as a promising domain for deep-tissue diagnostic, imaging and therapeutic applications, due to reduced photon scattering and increased tissue penetration depth enabled by low tissue autofluorescence in these wavelengths [6]. These NIR-emitting nanomaterials have several advantageous characteristics which make them suitable for biomedical applications, such as: (a) the ability to functionalize these materials with suitable polymer coatings and targeting agents for enhancing their biocompatibility, allowing for diverse applications in non-invasive detection of cancer and infectious diseases, drug delivery, photothermal and photodynamic therapy, to name a few; (b) the ability to be combined with other agents such as chemotherapy drugs to enable multi-modal treatment for maximal therapeutic efficacy; and (c) the low systemic toxicity afforded by the use of optically active NIR photothermal agents coupled with safe, non-ionizing light sources for cyclic treatment regimens; causing minimal photo damage to healthy tissue [7–10].

2D molybdenum disulfide (MoS_2_) are popular transition metal dichalcogenides because of their robustness, and unique band gap structure and physical, chemical and optical properties. MoS_2_ nanosheets have been investigated in the field of nanomaterials for biosensing and cancer therapy applications, since they exhibit strong near-infrared region absorbance and high photothermal conversion efficiency, and they constitute promising cancer phototherapy agents, since they heat up upon laser irradiation [11,12]. Also, their large surface area provides enhanced drug loading capacity and sites for binding functional ligands such as adhesive polymer chains (i.e. chitosan), and inorganic compounds such as Ag nanoparticles for enhanced antibacterial activity on medical devices or superparamagnetic iron oxide nanoparticles as a theranostic agent to facilitate bioimaging via NMR or MRI, and others [13–15].

In this study, a photo-activatable microparticle was developed for localized, pulsatile delivery of anticancer drugs into the tumor environment with simultaneous thermal ablation, by applying controlled ON-OFF thermal cycles using NIR laser irradiation. The microparticles were fabricated using a poly(caprolactone) (PCL) matrix, containing 2D MoS_2_ nanosheets as the NIR-responsive photothermal agent, and doxorubicin or violacein, as hydrophilic and hydrophobic model anticancer drugs, respectively. Cubic microparticles of 200 μm size were fabricated by casting a polymer-MoS_2_-drug film on polydimethylsiloxane (PDMS) mold with loading efficiency of 2 μg MoS_2_ per particle, and 0.2-8 μg of drug per particle. A cytotoxicity assay was performed to test the efficacy of the loaded microparticles in reducing the viability of 4T1 cell culture, using the ON-OFF laser switch mechanism. To select a suitable laser power and duration of treatment for *in vivo* studies, a machine learning algorithm based on a regression-based decision tree was developed to guide the design space for laser application, arriving at the optimal treatment conditions: 0.4 W/cm^2^ of laser power at 808 nm, for 3 cumulative minutes of laser irradiation, to reach the target temperature of ≥50 °C, which activates PCL melting and subsequent drug release. The effect of pulsatile treatment was studied *in vivo* in a 4T1 subcutaneous mouse model of breast cancer, using this light-activatable ON-OFF switch mechanism, for up to 3 cycles of laser irradiation. As control groups, tumor only (no treatment), laser only (no drug or microparticle), drug only (no laser or microparticle), microparticles only (no drug or laser), and microparticles with laser (no drug) were studied. The cohorts receiving the photo-activated 3-cyclic treatment for both drugs showed increased median survival up to 40 days post tumor induction, compared to a median survival of 16 days for the control groups. Although one laser cycle was enough to trigger drug release, there was a significant shrinkage in the tumor volume noticed with 3 cycles, with eventual scarring and shedding of the tissue with no evidence of residual tumor at the primary site.

## RESULTS AND DISCUSSION

Herein we describe the design of a novel strategy for cancer treatment which combines chemotherapy, in terms of targeted and controlled drug release, and phototherapy. 2D MoS_2_ nanosheets were produced via the liquid exfoliation method that supports lithium intercalation (top-down method) [13]. In Figure 1a the E_1g_, E^1^, and A_1g_ are first order Raman active modes resulting from vibrational modes within the S-Mo-S layer, whereas the distinct phonon modes at J_1_, J_2_ and J_3_ comprise indication of the metallic phase of the 2D MoS_2_. Figure 1b represents a TEM image of the MoS_2_ nanosheets in the range of 100-200 μm size, while Figure 1c demonstrated the elemental mapping for molybdenum and sulfur of the synthesized nanosheets based on EDS. Furthermore, in Figure S1 the cytotoxicity of MoS_2_ nanosheets was examined demonstrating that in the presence of MoS_2_ cell growth of 4T1 cells was not affected for concentrations as high as 200 μg/mL.

**Figure 1.**
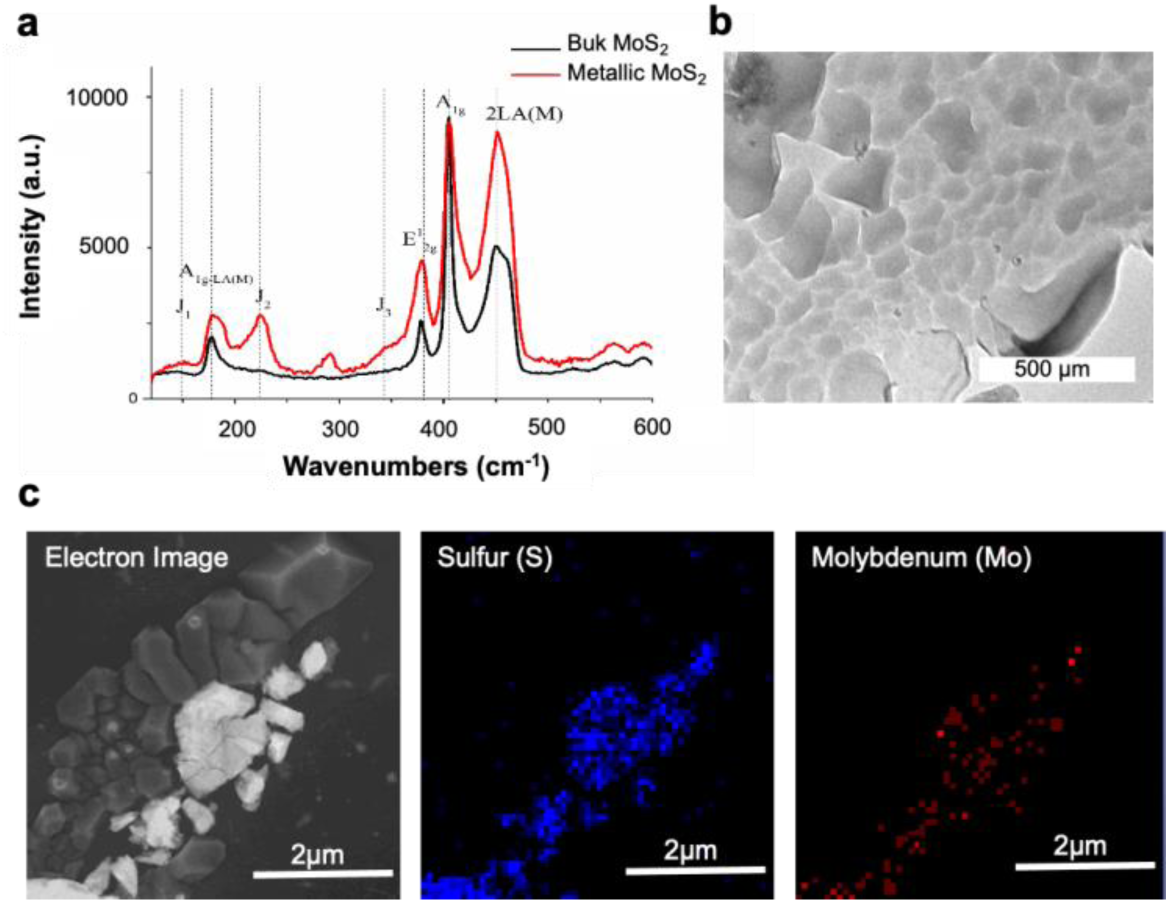
a) Raman spectra of bulk MoS_2_ versus metallic 2D MoS_2_ nanosheets, b) TEM imaging of 2D MoS_2_ nanosheets, c) EDS imaging of 2D MoS_2_ nanosheets; detection of molybdenum and sulfur.

As a following step cubic microparticles with 200 μm width were fabricated with the film casting technique. Briefly PCL, MoS_2_ nanosheets and the anticancer drug doxorubicin or violacein were dissolved in acetone and then cast on a heated Teflon surface to form a film. Subsequently, the formed film was heated and pressed into a polydimethylsiloxane (PDMS) mold to form solid microparticles. With this method, microparticles can be made with fidelity and high throughput (over 300 microparticles per array, 20 arrays per glass slide). Once the particles are formed, they get attached on the glass slide on which they are pressed, and they can be easily removed and collected in a tube by scraping off the surface of the glass slide. In Figure 2a formed particles are shown on a glass slide either loaded with doxorubicin or violacein. Violacein, an indolocarbazole compound, is a bacterial pigment with antimicrobial and anticancer properties against various cancer cell lines, such as breast cancer cells and uveal melanoma cell lines [16–20], however it has not been studied extensively since its commercial production is hindered by the low fermentation production yields. Nevertheless, violacein production is being subjected to synthetic biology routes that could potentially revolutionize its future use as drug scaffold [21]. Violacein was produced from *Janthinobacterium lividum* in a fed batch bioreactor and used after semi-purification. In Figure S2 the effect of different concentrations of the hydrophilic drug doxorubicin and hydrophobic violacein in semi-pure or pure form on 4T1 cell growth is presented, where it is evident that doxorubicin seems to be more potent comparing to semipure violacein, however pure violacein is even more effective in cell growth inhibition than doxorubicin. In Figure 2b SEM-EDS imaging shows the inorganic elemental composition of a formed PCL microparticle where the sulfur content is represented in pink and molybdenum in green. Furthermore, based on ICP analysis (Figure 2c) it was shown that the developed fabrication protocol results in approximately 1.8-2.0 μg of MoS_2_ loading per particle, while spectrophotometry after dissolving the particles in acetone and further dilution in water, demonstrated 0.2 μg and 8 μg loading of doxorubicin and violacein, respectively (Figure 2d). It is noteworthy that this fabrication method can facilitate the synthesis of different structures as shown in Figure S2, where microneedles and cylindrical microparticles are exhibited, along with needle performance when applied *ex vivo*.

**Figure 2.**
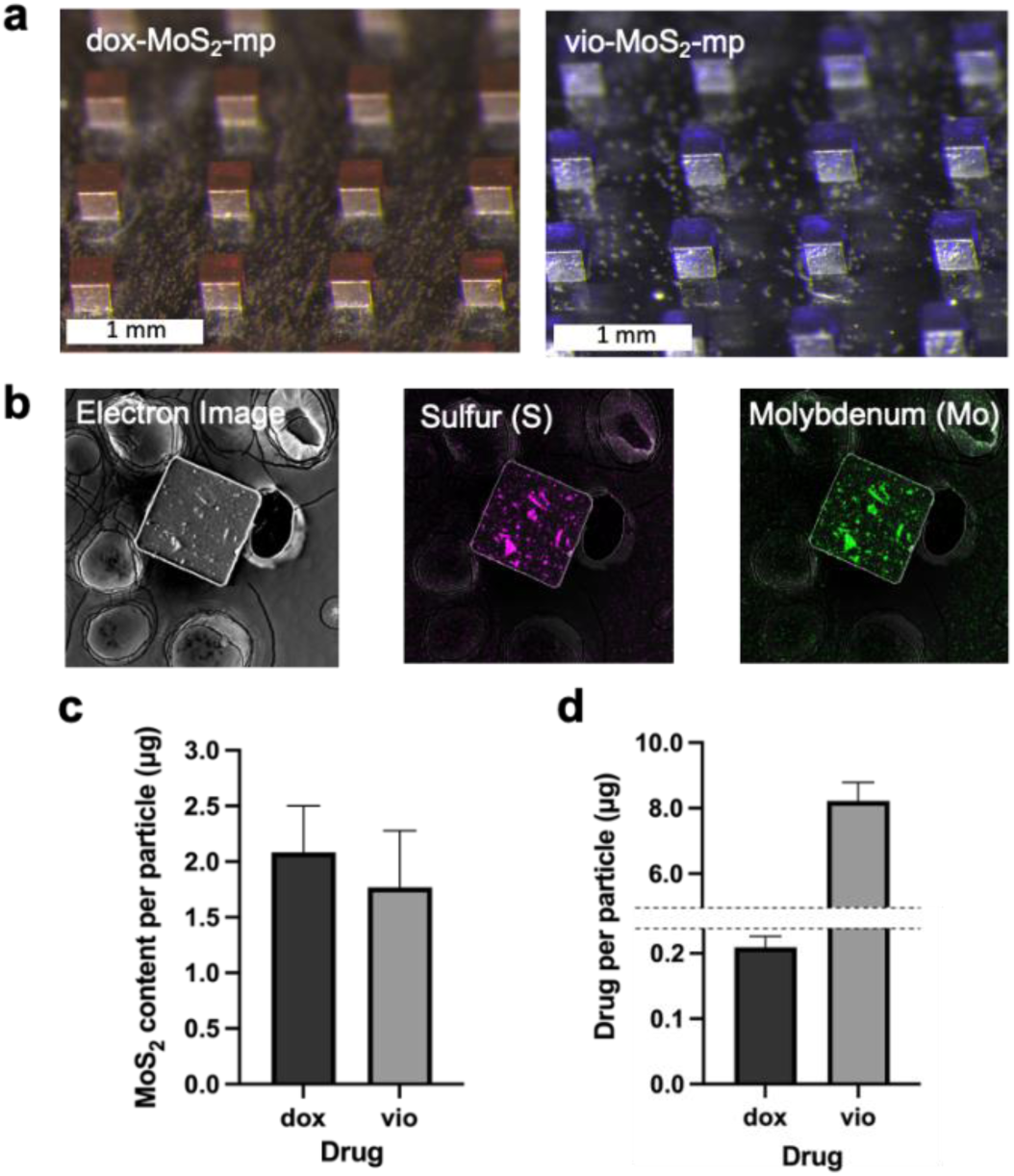
a) Optical imaging of fabricated drug and loaded microparticles loaded with MoS_2_ nanosheets and doxorubicin (dox-MoS_2_-mp) or violacein (vio-MoS_2_-mp), b) SEM-EDS imaging of one microparticles loaded with MoS_2_ nanosheets, c) Quantification of MoS_2_ content in the drug per particle based on ICP analysis, d) Quantification of loaded doxorubicin per particle based on fluorescence at 470 nm excitation and 560 nm emission wavelength, or violacein per particle based on absorbance at 575 nm wavelength.

Because of the MoS_2_ content the fabricated PCL microparticles are designed to warm up upon laser irradiation resulting to the PCL matrix melting and partial releasing of the embedded drug. To systematically understand the effect of the design variables on laser application and to decide on the operation conditions of the NIR laser, a design of experiments (DoE) study was carried out. The DoE matrix was constructed using Minitab software (Chicago, USA) based on an L54 Taguchi orthogonal array as shown in Table S1. Accordingly, 18 unique laser cycle conditions were investigated while each condition was repeated three times resulting in 54 experiments. Temperature was considered as the response while four design variables were considered including 1) the number of laser irradiation cycles, 2) duration of laser irradiation, 3) MoS_2_ amount and 4) laser power. Taguchi analysis was performed on these conditions to study the importance and ranking of design parameters on the resulting temperature upon laser application. DoE analysis demonstrated that MoS_2_ content, laser power, duration, and the number of cycles ranked as the most important to least important design parameter affecting the temperature, respectively.

As demonstrated in Figure 3a, all design parameters except the number of laser cycles, had a direct correlation with the temperature. Notably, the MoS_2_ mass demonstrated the highest increase in the mean temperature. Indicatively, increasing from 5 to 500 (μg) increased the mean temperature by approximately 30° C. Increasing laser power from 1 to 3 W, and laser duration from 1 to 3 min, led to a linear increase in temperature of approximately up to 15° C and 5° C, respectively. Conversely, increasing the number of laser cycles from 1 to 3 had a minimal effect on the resulting temperature, decreasing temperature by approximately 3° C.

**Figure 3.**
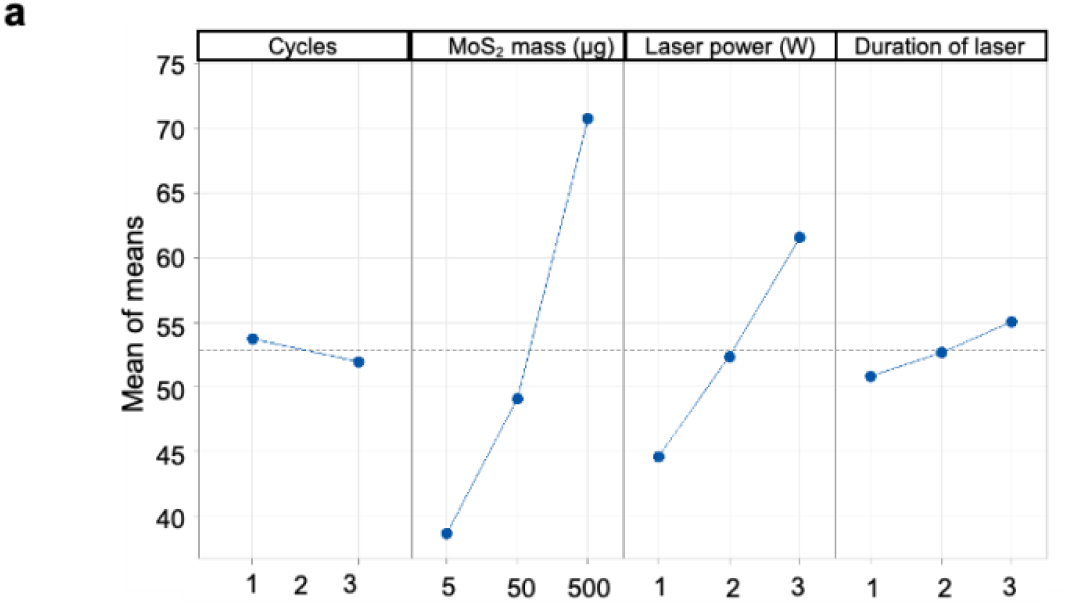
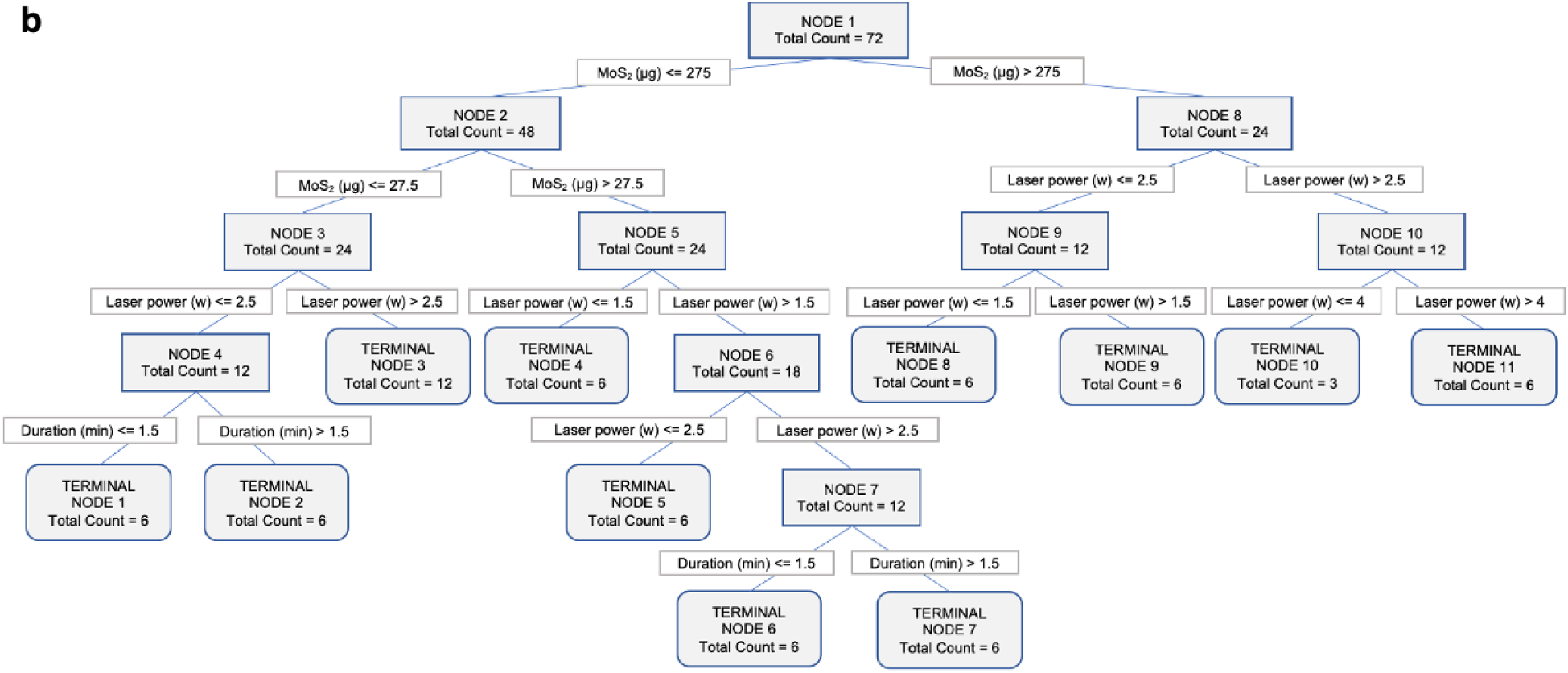
a) Study of the significance of the design parameters on the resulting temperature upon laser irradiation based on DoE analysis, b) An overview of the structure of the optimum decision tree developed to predict the resulting laser temperature as a function of MoS_2_ amount (in μg), laser power (in W), and laser duration (in min).

A linear regression model was also developed to predict the temperature for a given set of design variables. A dataset of 72 rows was used, 30.6% of which was used for testing and the rest used for training the model. The resulting training and testing *R*^2^ were found to be 83% and 80%, respectively. Such a model can be used as a preliminary design tool for similar studies to save costs associated with optimization experiments necessary to find the right laser power, duration and photothermal agent loading to achieve the desired temperature for effective PTT. The resulting regression model is as follows:

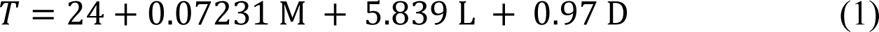

In which *T* is the resulting temperature *in vitro* in °C, *M* is the amount of MoS_2_ in μg, *L* is the laser power in W, and *D* is the laser duration in min.

As a following step, we sought to develop a Machine Learning based decision tree as a more sophisticated design framework to the regression model to systematically design the system for obtaining a certain temperature for laser PTT. This predictive decision tree framework is interpretable and can be a useful tool to achieve a specific target temperature upon application of laser for a specific treatment. Various hyperparameter optimizations were employed to optimize the structure of the decision tree. A plot of the structure of the optimal decision tree can be found in Figure 3b with the optimal hyperparameters provided in materials and methods section. A dataset containing 72 datapoints were utilized (Table S1), expanded for training and testing using 10-fold cross validation method. The optimal design resulted in an *R*^2^ percentage of 96.42%, and 93.73% for the training and testing datasets respectively, showing no signs of overfitting and highlighting excellent accuracy of the developed tree. The scatter plot of predicted *v*. actual value of temperature obtained by the decision tree is shown in Figure S3a showing very good prediction capability for both training and testing datasets. A ranking study of the design parameter was further carried out using the developed design tree. In agreement with the results from the DoE study, the design tree also highlighted MoS_2_ content, laser power, and laser duration as the most to least dominant design parameters on the laser temperature with relative importance values shown in Figure S3b while the number of cycles was considered insignificant.

To test the efficacy of the light-activatable system developed in this work, 25 particles loaded with dox or vio as the drug, and MoS_2_ nanosheets as the PTT agent, were incubated in 4T1 cell cultures and cytotoxicity tests were performed with and without laser treatment (Figure 4a). These conditions were also compared against (i) cells that received no treatment, (ii) cells that received free MoS_2_ nanosheets corresponding to 25 particles dose, with no laser treatment; (iii) cells that received free doxorubicin corresponding to 25 particles dose, with no laser treatment; (iv) cells that received free violacein corresponding to 25 particles dose, with no laser treatment; (v) cells that were incubated with 25 MoS_2_-microparticles, with no laser treatment and (vi) cells that were incubated with 25 MoS_2_-mps with laser treatment. In Figure 4b it is demonstrated that for cells receiving just MoS_2_ nanosheets in suspension or loaded in microparticles there was no growth inhibition when there was no laser treatment, suggesting that MoS_2_ does not induce cytotoxicity at the tested concentrations, without NIR light activation. Incubation with drugs violacein and doxorubicin at concentrations corresponding to the dose loaded in 25 microparticles inhibited cell growth up to 82%. Furthermore, cells that received vio-MoS_2_ microparticles without light activation did not affect cell growth, suggesting that the drug is well entrapped into the microparticles, considering that violacein in a hydrophobic compound, while dox-MoS_2_ microparticles induced ∼40% cell growth inhibition due to leakage of doxorubicin from the microparticles into the medium (color change of the medium was observed once particles were added), which may be attributed to the hydrophilic nature of doxorubicin. For groups MoS_2_-mp, vio-MoS_2_-mp and dox-MoS_2_-mp, upon laser irradiation for 3 min, microparticles’ melting was observed, and cell growth was inhibited by 70%, suggesting that activation of the particles induces pulsatile drug release. Laser irradiation of cells without MoS_2_ (suspension or particles) did not affect the cell growth, indicating that this type of irradiation can be specific and regioselective linked to the NIR phototherapeutic agent.

**Figure 4.**
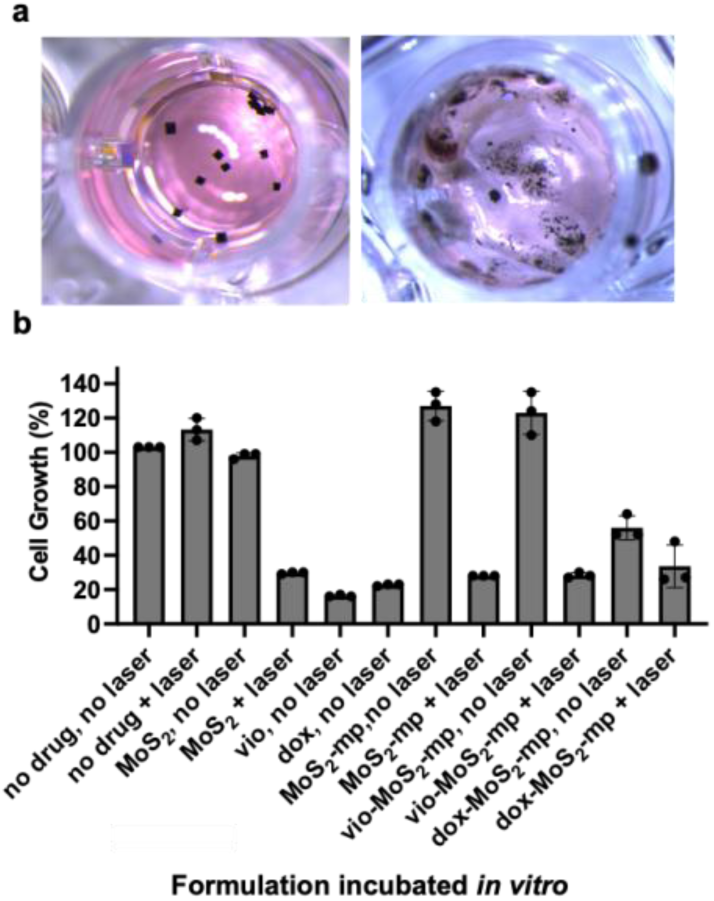
a) Incubation of vio-MoS_2_-particles in 4T1 cell culture before (left) and after (right) NIR laser irradiation, b) *In vitro* cytotoxicity assay in 4T1 cells under the various treatment conditions: no treatment, laser only (no drug or mps), free MoS_2_ (with or without laser), free drug (no laser), MoS_2_-mp (with or without laser), and vio/dox-MoS_2_-mp (with or without laser).

As a following step, the effect of combined photo- and chemotherapeutic treatment was studied *in vivo*, using a mouse model of 4T1 triple-negative breast cancer. Female Balb/c mice were injected in the right flank with a bolus of 2×10^6^ 4T1 cells in media, to induce subcutaneous tumor formation. Once mouse tumor size reached a threshold *V_T_* > 100 mm^3^, mice were divided into the various groups for treatment. The groups studied were: (i) control 1, no treatment; (ii) control 2, laser only with no MoS_2_; (iii) control 3, doxorubicin only, no laser; (iv) control 4, violacein only, no laser; (v) control 5, vio-MoS_2_-microparticles, no laser; (vi) MoS_2_-microparticles, with laser; (vii) dox-MoS_2_-microparticles with laser; (viii) vio-MoS_2_-microparticles with laser. Once the tumor size reached a threshold *V_T_* > 100 mm^3^ the mice received the 1^st^ NIR laser irradiation treatment. The temperature of the tumor was continuously monitored with a FLIR ONE thermal camera (Teledyne FLIR) and once temperature would reach 50 °C, laser treatment would be paused until temperature dropped to 39 °C, to avoid overheating at the tumor site. Total duration of laser irradiation per cycle was 3 cumulative minutes, as suggested based on the Machine Learning model. 3 NIR irradiation cycles were applied at the tumor site with a 3-day interval between cycles as suggested in the literature [22]. The laser ON-OFF and heating profile of the microparticles is presented in Figure S5.

Following tumor induction, mice were monitored regularly for body weight changes and tumor growth using physical measurements using a vernier caliper. Measuring the length, *L*, and the width, *W*, of the tumor along two orthogonal axes, the volume of the tumor was approximated using the ellipsoid formula, *V_T_* = 0.5 × *L* × *W*^2^. Representative images of the tumor growth for each group are shown in Figure 5, either for control groups right before euthanasia or for the treated group. vio-MoS_2_-mp over time.

**Figure 5.**
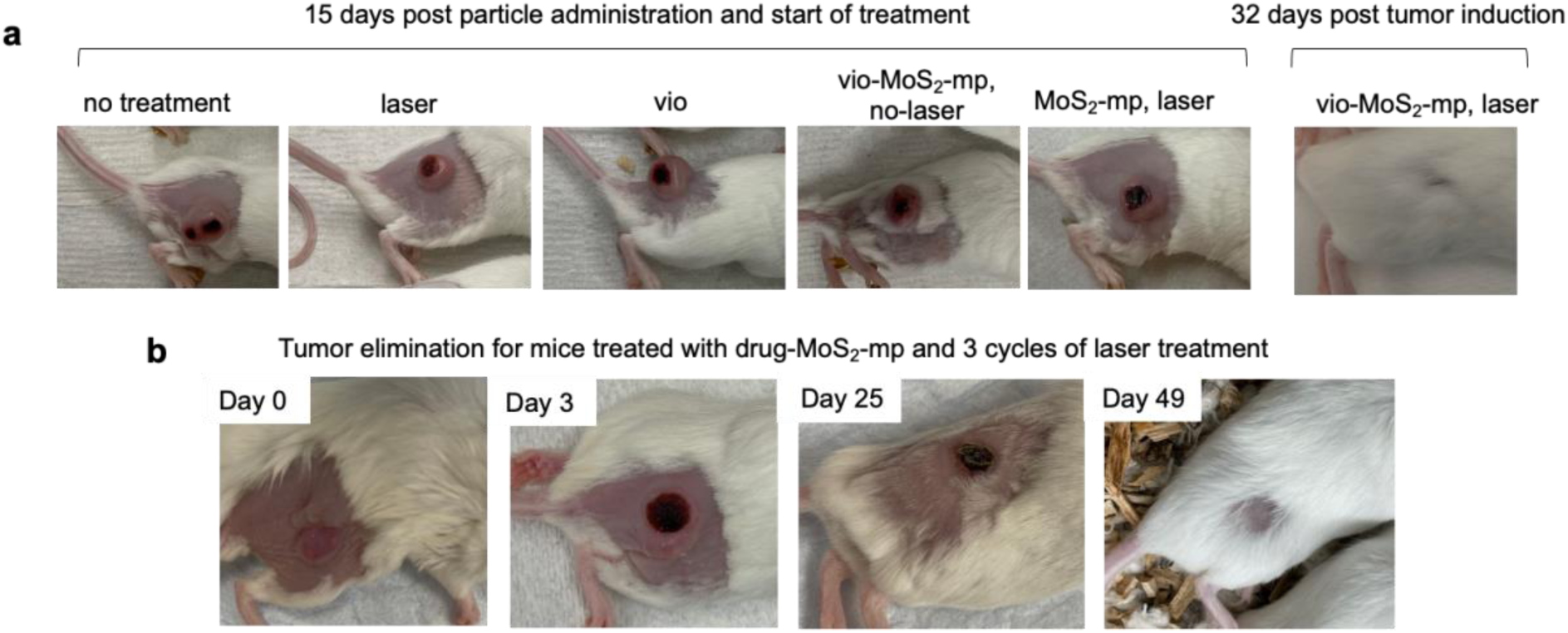
Representative images of a) mice from each group studied 15 or 32 days post treatment initiation, b) mice treated with vio-MoS_2_-microparticles and 3 cycles of laser irradiation over time.

The impact of the various treatment conditions on the mouse health was investigated using body weight as a metric, shown in Figure S6. At the start of the treatment (*t* = 0), mouse body weights were clustered in a narrow band in the range of 15-20 g. Over the duration of the treatment (or until the point of euthanasia), there were no significant differences in body weight observed among the various groups. Importantly, it should be noted that the optimized treatment groups, combining pulsatile PTT with on-demand drug delivery (vio-MoS_2_-mp or dox-MoS_2_-mp) groups did not show any drop in body weight, suggesting minimal adverse systemic toxicity of this treatment modality. The day of the first treatment cycle (laser only, drug only, or microparticles with drug and laser) is denoted as *t* = 0. The three laser cycles for NIR PTT are indicated on each plot as the black arrow with the red lightning bolt symbol, on *t* = 0, 3 and 6 days since the start of treatment.

The evolution of the tumor size for the various treatment groups, post start of treatment (*t* = 0), is plotted in Figure 6a. At the start of the treatment regimen (*t* = 0), mice in all groups had a very narrow range of tumor sizes, in the 100 mm^3^ range. For the no treatment, laser only, or drug only groups, there was a monotonic increase in the tumor sizes observed, with no appreciable shrinkage in the tumor volume. For the MoS_2_-mp with laser group (black line), there was an immediate decrease in the tumor volume after the first laser irradiation cycle, which was suppressed for the duration of the pulsatile NIR PTT (up to *t* = 6 days). However, the tumors started growing in volume almost immediately after the discontinuation of treatment, ie. after the completion of 3 laser cycles of PTT. This observation may explain the somewhat improved survival of the MoS_2_-mp group in the K-M curves in Figure 6b, but not quite as big an increase in the survival as the vio-MoS_2_-mp and dox-MoS_2_-mp groups. For the latter two treatment groups, the tumor sizes showed considerable shrinkage after the first cycle of treatment. More importantly, it was observed that after 3 cycles of combined PTT and synergistic pulsatile drug delivery, there was permanent shrinkage of the tumor, and no rebound in the tumor volume after the discontinuation of treatment. As shown in the representative images in Figure 5, with eventual scar formation and shedding of the tumor, there was no measurable residue of the tumor at the primary site. This dramatic inhibition of the tumor volume can explain the large increase in the median survival for these groups, observed in Figure 6b as noted above. Using pairwise *t*-tests, the differences in the measured tumor volume were found to be strongly significant for the vio-MoS_2_-mp / dox-MoS_2_-mp versus the control groups (no treatment, laser only, or drug only). While vio-MoS_2_-mp versus MoS_2_-mp was strongly significant, there was only weak significance in the difference between dox-MoS_2_-mp versus MoS_2_-mp, which could potentially be attributed to the initial sharp drop in the tumor size for the PTT-only MoS_2_-mp group, followed by a rebound in the tumor growth kinetics. Among the numerous control groups, the differences in the measured tumor volume were found to be non-significant; indicating that either laser PTT alone, drug release alone, or microparticles alone (without laser-activated synergistic PTT and chemotherapy) were insufficient to cause any meaningful decrease in the tumor burden.

**Figure 6.**
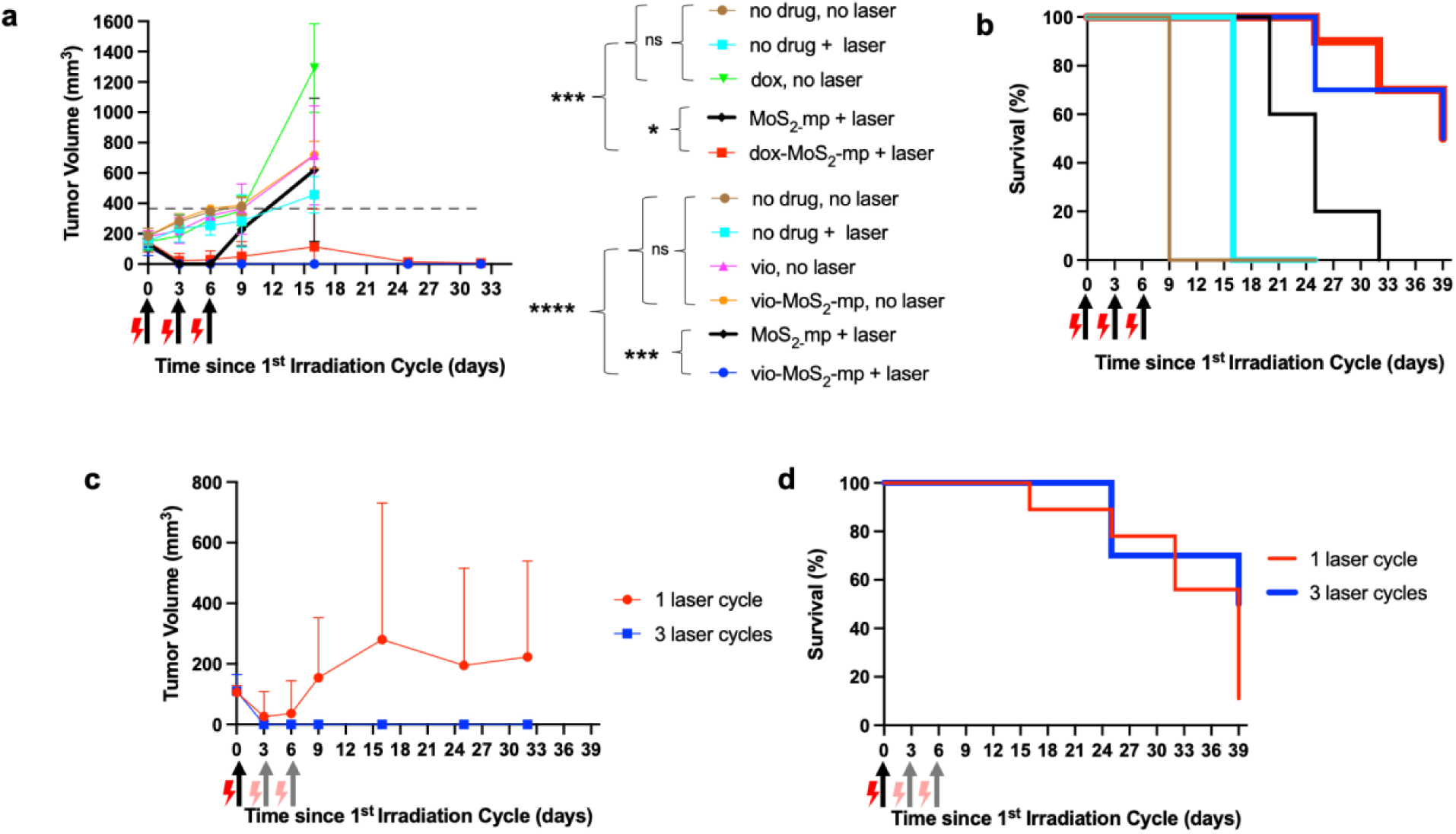
a) Evolution of tumor volume for control and treated groups since NIR irradiation treatment started as a function of time (dashed line shows the threshold criterion for euthanasia), reported as mean ± s.d. (*n* = 5 for the Control groups, *n* = 10 for the dox/vio-MoS_2_-mp groups). Student’s *t*-test used for pairwise significance, ɑ = 0.05. b) Kaplan-Meier survival curves for control and treated groups since NIR irradiation treatment started as a function of time, c) Evolution of tumor volume as a function of time, for vio-MoS_2_-mp, comparing 1- versus 3- cyclic laser treatment, reported as mean ± s.d. (*n* = 5). d) Kaplan-Meier survival curves as a function of time, for vio-MoS_2_-mp, comparing 1- versus 3- cyclic laser treatment.

The Kaplan-Meier survival curves for these different treatment conditions are plotted in Figure 6b. The no treatment and various control groups showed a sharp drop-off in the survival at *t* = 16 days, since these groups had to be euthanized because of the tumor size exceeded 9 mm (according to institutional policy, the Laboratory Animal Users’ Handbook of the Division of Comparative Medicine at MIT defines tumor size of 10 mm in diameter as criterion for euthanasia). While MoS_2_-mp with laser (phototherapy alone, no drug, black line) showed some improvement in the survival to *t* = 24 days, the strongest benefit was observed for the case of synergistic pulsatile on-demand chemotherapy, with ON-OFF phototherapy. As shown in the figure, for both the vio- MoS_2_-mp and dox-MoS_2_-mp cases (blue and red lines, respectively), compared to the untreated or drug only control groups, there was a significant increase in the median survival to *t* = 40 days post start of treatment.

Finally, the effect of the number of laser cycles on pulsatile, ON-OFF synergistic PTT and combined chemotherapy was studied for the vio-MoS_2_-mp treatment group, as shown in Figure 6c and Figure 6d. While for 3-cycle treatment (blue curve) the tumor volume is decreasing and staying low throughout the course of the treatment and beyond, interestingly, the 1-cycle treatment is more similar in nature to the MoS_2_-mp with laser (PTT only, black curve) in the sense that there is a moderate drop in the tumor volume in the first few days post therapy following a sharp rebound and rapid growth in the tumor. In terms of overall survival, by the end of the study only 10% survival is noted for the single cycle therapy, compared to 50% for the 3-cycle treatment. These results highlight the importance of multiple pulsatile PTT and on-demand controlled drug release cycles for better tumor management, compared to a single-dose phototherapy or single release drug delivery.

Aiming to demonstrate retention of the light-activatable microparticles at the tumor site with no systemic trafficking, vio-MoS_2_-microparticles were radiolabeled with ^89^Zr, and after intratumoral administration, PET imaging was performed before the start of each laser irradiation treatment (3 cycles of laser irradiation, with 3-day intervals between cycles) and in addition 3 days post the last (3^rd^) irradiation cycle. PET imaging exhibited retention of the radiolabeled particles at the tumor site for up to 12 days of imaging (Figure 7a). Furthermore, tissue from the tumor site was retrieved from mice there were untreated or treated with 1 laser cycle or 3 laser cycles for histological analysis. The tissue samples were fixed in formalin and embedded in paraffin and stained with Ki67 to evaluate the tumor’s proliferative activity. Mice that received the vio-MoS_2_-microparticles after 1 or 3 laser irradiations presented at least 26 % reduction in positive staining. Finally, a biodistribution study was performed in mice that received the vio-MoS_2_-microparticles and underwent 3 laser cycles. In this group (5 mice) tumors were eliminated at the primary site. At *t* = 40 days post laser irradiation, the mice were euthanized, and their spleen, liver and kidneys were harvested and treated with Aqua Regia before ICP-OES analysis to quantify the amount of molybdenum retained in each organ. Considering that initially 25 µg of Mo was injected at the tumor site (∼2 µg MoS_2_ loading per microparticle, for 25 microparticles injected: the total injected dose is ∼ 50 µg of MoS_2_, or effectively 30 µg of Mo), ultimately less than 0.3% of the injected dose is retained in the body.

**Figure 7.**
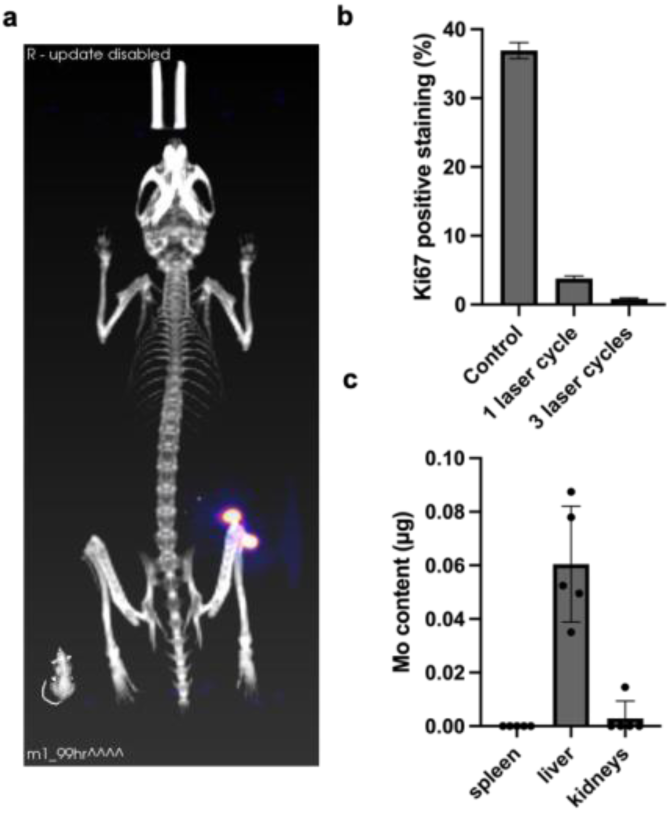
a) PET imaging of mouse lying in prone position with intratumorally injected ^89^Zr-labeled particles at the tumor site 3 days post 1^st^ laser irradiation cycle, b) Quantification of Ki67 staining of tumor site for untreated mice, mice treated with vio-MoS_2_-mp and 1 or 3 NIR irradiation cycles, c) Quantification of Mo content via ICP-OES in mice 40 days after treatment with vio-MoS_2_-mp and 3 NIR irradiation cycles, shown as mean ± s.d. (*n* = 5).

## CONCLUSIONS

To summarize, in this work, we have developed near-infrared responsive MoS_2_-based microparticles for on-demand, pulsatile phototherapy and drug delivery, using Machine Learning-enabled optimization to achieve the best therapeutic efficacy for our combination therapy. The efficacy of the system was demonstrated in a subcutaneous mouse model of triple-negative breast cancer. While other approaches to combined PTT with drug delivery have been previously reported in the literature, this work presents a paradigm-shifting advancement in three key ways: (1) The use of Machine Learning to tailor the right conditions of PTT, by enabling the informed choice of an optimal loading of MoS_2_, the laser power density, and duration of irradiation cycles resulting in maximal efficacy of tumor treatment; (2) The use of low-dose, localized chemotherapy, 5-200 μg of drug (compared to >1000 μg in the literature), coupled with on-demand pulsatile low-power laser PTT, 0.4 W/cm^2^ (compared to 5 W/cm^2^ in the literature), resulting in low systemic toxicity; (3) The ability to achieve complete eradication of the primary tumor site, with sustained tumor inhibition maintained up to 40 days after the completion of 3 cycles of treatment, resulting in enhanced median survival compared to the no treatment or monotherapy groups. Given these significant advancements, we strongly believe that upon further optimization of the microparticle loading capacity and clinical translation, our synergistic photo- and chemotherapeutic treatment approach has the potential to achieve better response outcomes for patients with breast cancer, melanoma, or similar subcutaneous tumors.

## MATERIALS AND METHODS

### Materials

PCL (MW = 70 kDa), n-butyllithium and MoS_2_ bulk were purchased from Sigma-Aldrich (St. Louis, MO, USA). Doxorubicin hydrochloride was obtained from TCI (Tokyo, Japan). Polydimethylsiloxane (PDMS, Sylgard 184) was provided by Dow Corning (Midland, MI).

### 2DMoS_2_ nanosheets synthesis and characterization

For the synthesis of MoS_2_ nanosheets 5 mL of n-butyllithium in hexane was added in 1mg of MoS_2_ bulk powder under the protection of nitrogen gas. After 2 days of intercalation, the MoS_2_ solution was washed with hexane. Raman analysis was performed on Renishaw Invia Reflex Micro Raman (Gloucestershire, United Kingdom) equipment. High resolution imaging with Transmission Electron Microscopy-Energy Dispersive X-ray Spectrometry was performed using an JEOL 2100 FEG microscope. All images were recorded on a Gatan 2kx2k UltraScan CCD camera. STEM imaging was done by HAADF (high-angle annular dark field) detector. X-Max 80 mm^2^ EDX (Oxford Instrument, UK) was used for chemical information of samples.

### Violacein production

Violacein from *Janthinobacterium lividum* was produced in a fed-batch bioreactor as described previously and it was used in semi-pure form [23,24].

### Fabrication of MoS_2_-based microstructures

#### Fabrication of cubic or cylindrical microparticles

133 mg of PCL, 40 mg MoS_2_ nanosheets and 50 mg of the anticancer drug doxorubicin or violacein were dissolved in 2 mL acetone and then cast on a heated Teflon surface to form a film. Subsequently, the formed film was heated and pressed into a polydimethylsiloxane (PDMS) mold to form solid microparticles. With this method microparticles can be made with fidelity and high throughput (over 300 microparticles per array, 20 arrays per glass slide). Once the particles are formed, they get attached on the glass slide on which they are pressed, and they can be easily removed and collected in a tube by scraping off the surface of the glass slide.

#### Fabrication of microneedle arrays

2.4 g of PCL, 55 mg of MoS_2_ nanosheets and 65 mg of the anticancer drug doxorubicin or violacein were dissolved in 9.6 mL acetone. For each microneedle patch (1×x1 cm^2^ footprint, 1.5mm needle height, 0.4mm needle width, 1mm spacing between needles, 1mm patch backing) 200 μL were dispensed on negative PDMS molds covered with filtration paper, following centrifugation for 10 min at 4200 rpm. This process results in forming drug-MoS_2_-PCL needle tips. The molds’ surface were wiped with acetone rinsed wipes and left to dry overnight at 60 °C and then to form the backing of the needles 200 μL of 20 wt.% poly(vinyl alcohol)(PVA, 31kDa)-poly(vinylpolypyrrolidone) (PVP, 10kDa) was dispensed on the molds and centrifuged at 4200 rpm for 5 min, following by addition of 200 μL and another 300 μL 30 min later. The patches were left to dry for 3 days at room temperature and then were removed from the molds and stored in a desiccator under vacuum.

### Machine learning algorithm for MoS_2_ NIR laser conditions

A classification and regression tree (CART) was designed in Minitab incorporating the data from 72 experiments related to application of laser. The structure of the decision tree was optimized to avoid overfitting while maximizing accuracy of the test set. The following parameters were used in the optimal version of the decision tree, number of terminal nodes of 11, minimum terminal node size of 6, node splitting method of least square error (LSE), model validation method of 10- fold cross validation.

### Optical imaging

The fabricated microstructures and the incubated cubic microparticles *in vitro* were imaged via a Leica DFC450 Optical microscope using with LAS V4.7 software.

### NIR Laser

A continuous wave, Class IV multimode laser at 808 nm was used (MDL-N-808 diode pumped solid state laser, Opto Engine LLC, Midvale, UT, USA). An achromatic beam expander (GBE05-B, Thorlabs, Newton, NJ, USA) was coupled to the optic fiber output of the laser, to achieve a 5x areal expansion, with an output beam aperture of 35 mm. An optical power meter (PM100D, Thorlabs) was used to measure the output laser power. Based on the Machine Learning algorithm the optimal power density was determined to be 0.4 W/cm^2^.

### 4T1cell culture

The mouse breast cancer cell line 4T1 (ATCC CRL-2539) was cultured in Dulbecco’s modified essential medium supplemented with antibiotics containing 100 U/mL penicillin, 100 mg/mL streptomycin, and 10% fetal bovine serum. The cells were then incubated in a humidified atmosphere containing 5% CO_2_ at 37 °C.

### Tumor induction and dosing of microparticles

Female Balb/c mice, 4-6 weeks old, were procured from Charles River (MA, USA). Mice were housed in cages with access to food and water *ad libitum*. After an acclimatization period of 1 week, mice were prepared for tumor injection by shaving and chemical depilatory treatment in the right upper flank region. 4T1 tumor cells were suspended in culture media, and injected in the right upper flank at a concentration of 2×10^6^ cells in 50 μL of medium.

Post tumor induction, the tumor volume was measured regularly (every 2-3 days) using orthogonal axis measurements using a vernier caliper. The tumor volume was approximated using the ellipsoid formula. Once the tumor volume was determined to be above a threshold of 100 mm^3^, the animals were ready for treatment. The mice were randomly divided into eight groups with 5 mice per group for the controls and 10 mice per group for the drug and MoS_2_ loaded particle groups that underwent NIR laser irradiation. For groups that received particles; 25 particles were administered intratumorally using an 18G perfusion needle to avoid clogging of the syringe. The particles were suspended in 1% carboxymethyl cellulose in PBS buffer.

### Quantification of Mo with Inductively Coupled Plasma

For quantification of Mo content in vitro the particles were dissolved in hydrochloric acid and diluted in water. For quantification of Mo content *in vivo*, mouse tissue and organs were dissolved in Aqua Regia medium (nitric acid: hydrochloric acid 1:3) and diluted in water. The samples were analyzed using Inductively Coupled Plasma - Optical Emission Spectrometer (ICP-OES) Agilent ICP-OES 5100 VDV (Santa Clara, CA, USA).

### SEM-EDS

SEM and EDS-SEM were performed using a JSM-5600LV SEM (JEOL, Tokyo, Japan) with an acceleration voltage of 5 to 10 kV. Samples were initially coated with a thin layer of Au/Pd using a Hummer 6.2 sputtering system (Anatech, Battle Creek, MI) and then imaged. EDS was performed in the dark field, and both SEM and EDS-SEM were performed at high vacuum settings.

### Radiolabeling of vio-MoS_2_-microparticles with ^89^Zr

Initially amine groups were applied on the surface of the PCL microparticles covalently through aminolysis; 30 microparticles are incubated in a vial and washed 3 times with 70 % ethanol. After drying at room temperature, to perform aminolysis at room conditions 2 mL of 2 % w/v 1, 6-hexanediamine in 2-propanol is added and the particles are incubated at 37°C for 24 h under stirring [25]. Then the particles are washed 3 times with water and dried at room conditions. The particles are then incubated with PBS buffer with adjusted pH at 8.0 using 1 M dipotassium phosphate with the chelator p-SCN-Bn-Deferoxamine (B-705, Macrocyclics, Plano, TX, USA) overnight at 4 ℃. The chelator is dissolved initially in DMSO at 5 mg/mL. In the vials that contain the dried particles post aminolysis, 1.6 mL of PBS pH 8.0 is added and 200 μL of the chelator stock. The particles are incubated at 4 °C for 24 h. The buffer is treated with Chelex 100 Resin (142-1253, BioRad) to scavenge free iron to prevent contamination of the chelator.

### Quantitative analysis of Ki-67 tumor staining

For Ki-67 staining tissue samples from the tumor site were retrieved fixed in formalin and embedded in paraffin. The samples were then cut into 10 sections of 5μm thickness prior to staining. 2D image analysis of tissue cross-sections from three groups (control, laser 1 cycle, laser 3 cycles) with n = 20, 20, 40 respectively, stained with KI-67 was performed using the open source software QuPath (UK) and ImageJ (USA). Pre-processing steps were applied on each image to prepare them for subsequent analysis using an image analysis thresholder. The thresholder was created to isolate tissue samples from the image background and designate these sections as regions of interest in the software. For the Ki67 stain, we processed images using a positive cell detection function to identify the number and percentage of positively stained cells (brown). After trial and error with manipulating the automatic settings given by the software, the following settings were then used for the cell detection functionality: detection image, optical density sum; requested pixel size = 1 µm; background radius = 8 µm; median filter radius = 1 µm; sigma = 2 µm; minimum cell area = 11 µm^2^; maximum cell area = 300 µm^2^; threshold = 0.2; maximum background intensity = 2. Moreover, the following settings were utilized to detect Ki-67 positive staining in the detected cells: score compartment, Nucleus: DAB OD mean; Threshold 1, 0.5. This function was run on all regions of interest and was able to output the percentage of cells that were positively stained for KI-67 by counting the relevant cell nuclei.

## ASSOCIATED CONTENT

### Supporting Information

The following files are available free of charge.

-Real-time thermographic video of microparticles being irradiated after intratumoral injection *in vivo* during first irradiation cycle (AVI).

-Supplementary material including (PDF):

- Cytotoxicity studies for MoS_2_ nanosheets, violacein and doxorubicin at different concentrations
- Imaging of other produced microstructures with embedded drug and other MoS_2_ nanosheets
- Scatter plot of fitted versus actual laser temperature obtained predicted by the developed decision tree algorithm, and relative importance of design parameters used in the decision tree to predict the temperature value attained for a given laser application.
- Optical imaging of cross-section of pig tissue where dox-MoS_2_-mpicroparticles were injected subcutaneously, before and after 3min NIR laser irradiation.
- Laser irradiation and heating profile per cycle, *in vivo*.
- Body weight for control and treated mice groups since NIR irradiation treatment started as a function of time.

### Ethics statement

All animal handling and procedures were done in compliance with the Institutional Animal Care and Use Committee protocols. Procedures were pre-approved by MIT’s Committee on Animal Care.

### Statistical analysis

Statistics were performed in GraphPad Prism using Student’s *t*-test for pairwise comparisons at a significance level of α = 0.05. All *in vitro* experiments were performed in experimental triplicate or quintuplicate. All *in vivo* experiments in mice were performed with 5 or 10 experimental replicates (2 independent experiments 5 mice each). Data are reported in the text as mean ± s.d.

### Notes

The authors declare no competing financial interest.

### Author Contributions

The manuscript was written through contributions of all authors. All authors have given approval to the final version of the manuscript.

MK: Design of experiments, conceptualization, and fabrication of photo-activatable system, animal handling, data analysis, writing manuscript.

NB: Design of experiments, optimization of NIR laser beam area, optical power and time of laser irradiations for *in vivo* tumor studies, developing chelation protocol for conjugating ^89^Zr to the

PCL matrix for PET imaging, monitoring animal cohorts over long term (up to *t* = 60 days) post treatment, writing manuscript

MS: Machine learning application, DoE analysis, EDS-SEM imaging

SA: ICP analysis

WR: Excision of tumors and organs on animal terminals, histology evaluation

AP: Induction of tumors in animals

D de F: Development of method and processing of Ki67 stained histology slides HM, VS, NH: Handling of radio-labeled materials and PET imaging

JH: Graphical abstract

AB, AJ, RL: Design and conceptualization of the project, data analyses, manuscript review

## ACKNOWLEDGMENTS

This work was supported by the Bodossaki Foundation (Athens, Greece) with a postdoctoral scholarship to MK. Furthermore, this work was supported by the Onassis Foundation under the Special Grant & Support Program for Scholars’ Association Members (Grant No. R ZQ 002-1/2020-2021). N.M.B. acknowledges funding support through the Mazumdar-Shaw International Oncology Fellowship. W.TR. is funded by National Cancer Institute fellowship 5T32CA009216-40. We thank the Koch Institute’s Robert A. Swanson (1969) Biotechnology Center (RRID: SCR_018674) for technical support, specifically the Histology core, the Nanotechnology Materials core, the BioMicroCenter, and the Animal Imaging and Preclinical Testing facilities. This work was supported in part by the Koch Institute Support (core) Grant P30-CA14051 from the National Cancer Institute. We would like to thank MIT animal facilities for the *in vivo* studies in mice.

## ABBREVIATIONS

MoS_2_: molybdenum disulfide
PCL: poly(caprolactone)
PDMS: poly(dimethylsiloxane)
vio: violacein
dox: doxorubicin
Ki-67: protein encoded by the MKI67 gene, marker of cell proliferation
MoS_2_-mp: microparticles loaded with MoS_2_
dox-MoS_2_-mp / vio-MoS_2_-mp: microparticles loaded with MoS_2_ and drug (either doxorubicin or violacein)
NIR: near infrared
DoE: design of experiments
ICP-OES: inductively coupled plasma optical emission spectrometer
PET: positron emission tomography
PTT: photothermal therapy
SEM-EDS: scanning electron microscope with energy-dispersive X-ray spectroscopy

