## Supplementary Information for "A Machine Learning-optimized system for on demand, pulsatile, photo- and chemo-therapeutic treatment using near-infrared responsive MoS_2_-based microparticles in a breast cancer model"

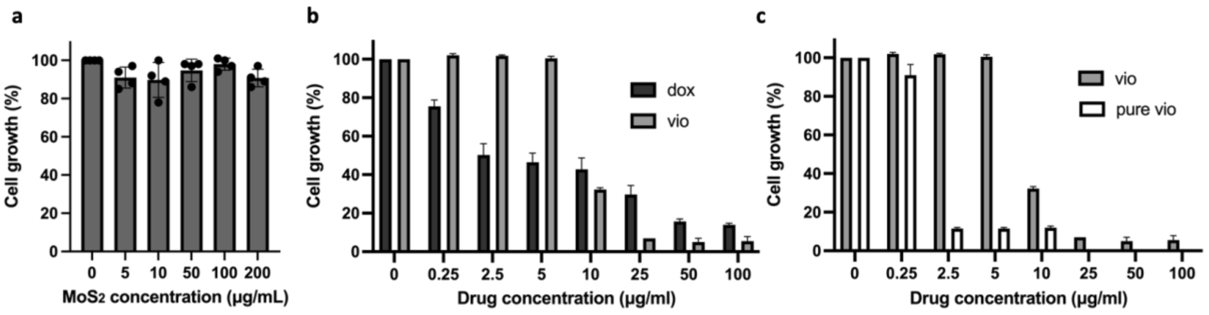


**Fig. S1:** Cytotoxicity study in 4T1 cells for a) different concentrations of $MoS_2$ nanosheets, b) drugs doxorubicin versus semi-pure violacein, c) violacein in semi-pure versus pure form.

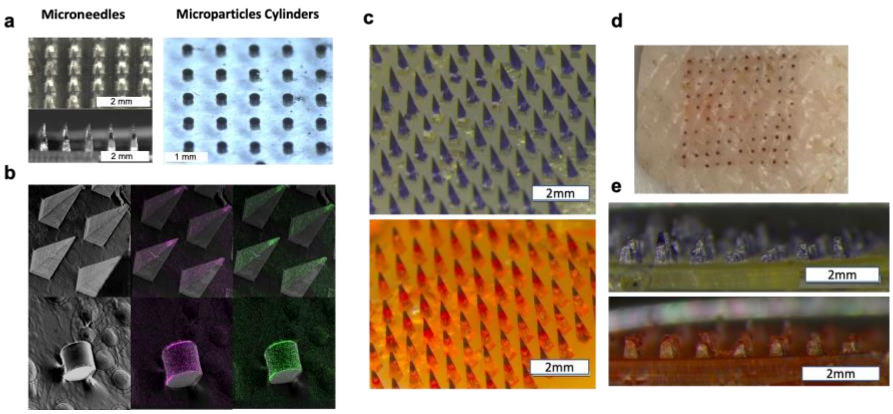


**Fig. S2**: a) Optical imaging and b) SEM-EDS imaging of microneedles and cylindrical microparticles with embedded $MoS_2$ nanosheets, c) Optical imaging of microneedle patches with with violacein or doxorubicin + $MoS_2$ nanosheets embedded in PCL tips, d) Needles' footprint post application of patch *ex vivo* on pig cadaver, e) Needle dissolution after 10 min application *ex vivo* on pig skin.

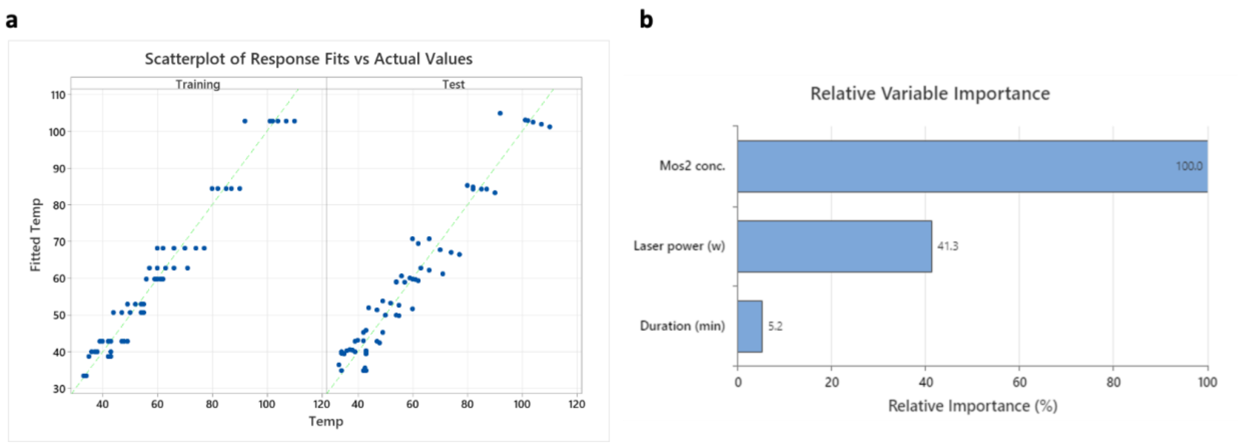


**Fig. S3:** a) Scatter plot of fitted versus actual laser temperature obtained predicted by the developed decision tree algorithm. b) Relative importance of design parameters used in the decision tree to predict the temperature value for a laser application.

a b c

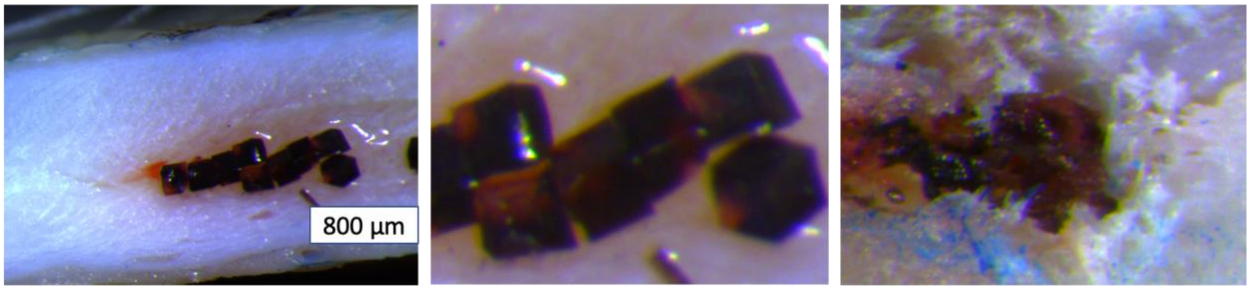


**Fig. S4:** a) Optical imaging of cross-section of pig tissue where dox-$MoS_2$-mp were injected subcutaneously, b) Zoom-in on particles injected before NIR laser irradiation, c) Zoom-in on particles after 3 min NIR laser irradiation.

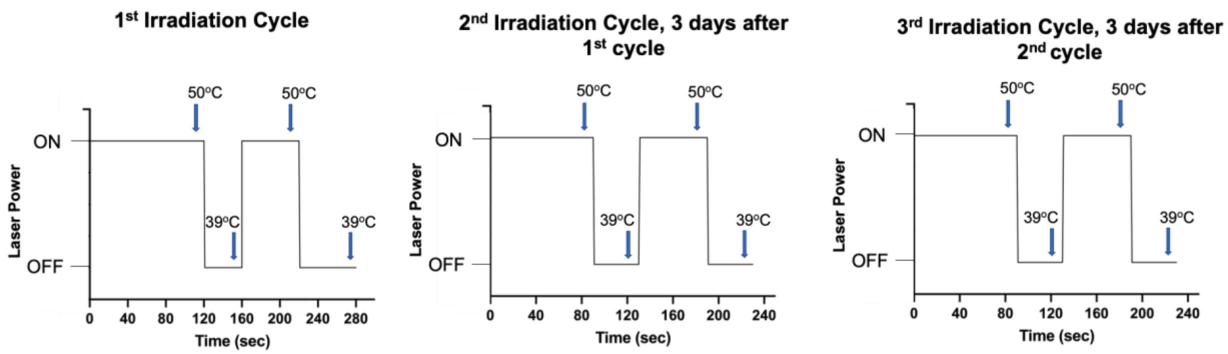


**Fig. S5:** Laser irradiation and heating profile per cycle, *in vivo*.

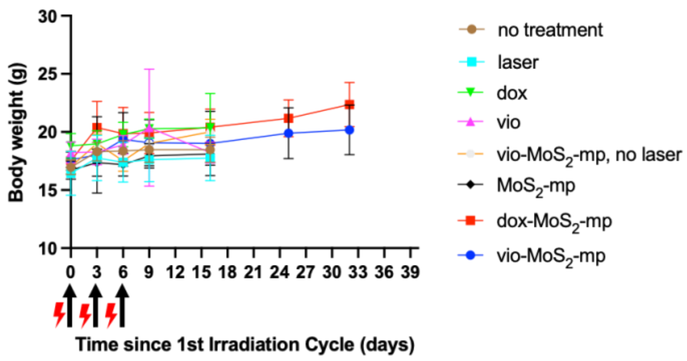


**Fig. S6:** Body weight for control and treated groups since NIR irradiation treatment started as a function of time.
